# OpenAc4C: A gateway to decode the landscape, regulation and pathogenesis of N4-acetylcytidine (ac^4^C) epitranscriptome

**DOI:** 10.64898/2026.04.01.714364

**Authors:** Gang Tu, Yigan Zhang, Xuan Wang, Jishuai Zhang, An Zhu, Kunqi Chen, Zhixing Wu, Zekai Wu, Yue Wang, Jingxian Zhou, Zhen Wei, Guifang Jia, Jia Meng, Daniel J. Rigden, Bowen Song

**Affiliations:** Department of Public Health, School of Medicine, Nanjing University of Chinese Medicine, Nanjing 210023, China; Institute of Biomedical Research, Regulatory Mechanism and Targeted Therapy for Liver Cancer Shiyan Key Laboratory, Hubei Provincial Clinical Research Center for Precise Diagnosis and Treatment of Liver Cancer, Taihe Hospital, Hubei University of Medicine, Shiyan 442000, China; Department of Biosciences and Bioinformatics, Center for Intelligent RNA Therapeutics, Suzhou Key Laboratory of Cancer Biology and Chronic Disease, School of Science, XJTLU Entrepreneur College, Xi’an Jiaotong-Liverpool University, Suzhou 215123, China; Institute of Systems, Molecular and Integrative Biology, University of Liverpool, Liverpool L7 8TX, United Kingdom; Synthetic and Functional Biomolecules Center, Beijing National Laboratory for Molecular Sciences, Key Laboratory of Bioorganic Chemistry and Molecular Engineering of Ministry of Education, College of Chemistry and Molecular Engineering, Peking University, Beijing 100871, China; Key Laboratory of Ministry of Education for Gastrointestinal Cancer, School of Basic Medical Sciences, Fujian Medical University, Fuzhou 350004, China; Jiangsu Key Laboratory for Functional Substance of Chinese Medicine, School of Pharmacy, Nanjing University of Chinese Medicine, Nanjing 210023, China; Sino-French Hoffmann Institute, School of Basic Medical Sciences, Guangzhou Medical University, Guangzhou, Guangdong 511436, China

**Keywords:** N4-acetylcytidine (ac^4^C), Epitranscriptome, Deep learning, Disease-variant, ac4C Finder

## Abstract

N4-acetylcytidine (ac^4^C) is an ancient and highly conserved chemical marker found in all domains of life. Recent advancements in sequencing techniques have enabled the functional analysis of ac^4^C occurrence by accurately capturing its locations and levels, shedding light on its significant regulatory potential and emerging role in diseases. The OpenAc4C, the first comprehensive knowledgebase exclusively designed for unraveling the ac^4^C epitranscriptome across diverse species, spanning vertebrates, mammals, insects, fungi, plants, bacteria, archaea, and viruses. By mining a large array of ac^4^C epitranscriptome datasets with deep learning-based pipelines, OpenAc4C features a collection of 536,745 ac^4^C sites identified from four distinct next-generation sequencing (NGS)-based techniques, alongside novel insights from Oxford Nanopore direct RNA sequencing (ONT)-based samples, encompassing a total of 33 species. Beyond the ac^4^C landscape, a total of 536,986 ac^4^C-affecting variants were identified in seven species. Among them, 4,766 pathogenic ac^4^C-SNPs may drive ac^4^C dysregulation with implications for disease pathogenesis. In addition, OpenAc4C offers a user-friendly graphical interface and a web-based analysis platform for comprehensive querying and interactive exploration of the database collections. Together, OpenAc4C will serve as a valuable integrated resource to facilitate studies of ac^4^C modification. It is freely accessible at: www.rnamd.org/ac4cportal.

## Introduction

The term ‘epitranscriptomics’ refers to the study of a broad range of chemical modifications that naturally occur on various RNA types [1, 2], dynamically modulating nearly every stage of RNA fate such as mRNA stability, splicing event, transport, translation and degradation [3–6]. Among the over 170 types of modified residues, N4-acetylcytidine (ac^4^C) is an ancient and highly conserved chemical marker initially detected in the bacterial tRNA^Met^ anticodon [7]. It was subsequently reported in tRNA, rRNA and mRNA as a universally conserved modification in all domains of life. In eukaryotic RNA, ac^4^C was found to be the sole acetylation mechanism catalyzed by the acetyltransferases N-acetyltransferase 10 (NAT10) and Kre33 in human and yeast, respectively [8], modulating diverse cellular processes such as gene expression regulation [9], protein translation efficiency [10], ribosome maturation [11], and RNA stability [12, 13]. Notably, recent studies have also suggested that ac^4^C is involved in the occurrence and progression of various diseases including cancer [14–16]. In cervical cancer, NAT10/ac^4^C/FOXP1 axis activity enhances the immunosuppressive properties of tumor-infiltrating regulatory T cells (Tregs) by promoting glycolysis in the continuously lactic acid-enriched tumor microenvironment (TME) [17]. Meanwhile, ac^4^C modifies and stabilizes the mRNA of Ferroptosis suppressor protein 1 (FSP1) and thus suppresses ferroptosis, which in turn contributes to the progression of colon cancer [18]. Taken together, these studies have greatly expanded our understanding on ac^4^C acetylation and implied its significant regulatory potentials.

To date, a number of high-throughput sequencing techniques have been proposed for the global mapping of ac^4^C modification. The first transcriptome-wide approach to investigate ac^4^C localization was introduced in 2018 using RNA immunoprecipitation with anti-ac^4^C antibody (ac4C-RIP-seq or acRIP-seq) [19]. This technique generates ac^4^C-enriched regions (peaks) with a resolution of approximately 150 bp and has proven invaluable in profiling ac^4^C level across multiple species, including human [19], mouse [20], rat [21], cellular mRNAs of KSHV transcripts [12] and others [22]. In addition to acRIP-seq, photo-assisted (PA)-ac4C-seq employs PA crosslinking of ac^4^C-specific antibodies to 4-thiouridine (4SU)-labeled RNAs and has been used for mapping ac^4^C residues in HIV-1 mRNA with improved resolution [23]. Recent advances such as ac4C-seq [24], RedaC:T-seq [25], and RetraC:T-seq [26] offer the precise detection of ac^4^C modification sites at base-resolution level. It is worth noting that detection sensitivity varies across different profiling techniques. The base-resolution technique ac4C-seq, pursued in HEK-293T, identified ac^4^C sites in mRNA only under specific condition of NAT10 overexpression [24], while methods of PA-ac4C-seq [23], acRIP-seq [19], RedaC:T-seq [25], RetraC:T-seq [26], and liquid chromatography-tandem mass spectrometry (LC-MS/MS) consistently confirmed the presence of ac^4^C in mRNA [2, 19, 27]. In addition to the widely applied next-generation sequencing (NGS)-based methodologies, the presence of RNA modified residues can be simultaneously detected as characteristic current signals using Nanopore direct RNA sequencing [28], proposed by the third generation sequencing platform of Oxford Nanopore Technology (ONT). Various tools have been developed for profiling specific modification types such as m^6^A (m6Anet and MINES) [29, 30], pseudouridine (Ψ) and m^1^Ψ (nanoPsu and NanoMUD) [31, 32], or ELIGOS [33] for identification of mixed (unlabeled) modified residues from nanopore direct sequencing data. Of note, software tools such as ELIGOS and Tomobo currently lack the capability to differentiate specific modification types from common base-calling errors [33]. These unknown types of modified residues could potentially be further labeled through mapping with NGS-based modification sites or deep learning methods [34].

As our understanding of RNA modification continues to expand, bioinformatics efforts have been made to annotate, interpret, and share the rapidly generated epitranscriptomic datasets. These databases encompass a broad array of modification types, each with a distinct functional emphasis: MODOMICS for querying RNA modification (RM) pathways with new feature on data annotation system [35–37], ConsRM for quantitively measuring the conservation degree of m^6^A methylation [38–40], RMBase v3.0 for revealing the map of RMs in multiple species [41], RMVar and RMDisease for unveiling RM-affected genetic variants [42, 43], and RM2Target for collecting RM-dependent enzymes [44]. However, epitranscriptome databases such as RMBase and RMDisease were developed to encompass extensive collections of diverse RNA modification types rather than focusing on a specific one [45, 46], as a result, they offer only very limited information on ac^4^C acetylation. To the best of our knowledge, none of the current resources have been designed to systemically decipher the various aspects of ac^4^C biology, particularly considering its significant regulatory roles and the massive datasets revealed recently.

In this study, we presented OpenAc4C, the first integrated resource to decode the transcriptome landscape, regulatory mechanisms, and disease pathogenesis of ac^4^C modification across diverse species spanning vertebrates, mammals, insects, fungi, plants, bacteria, archaea, and viruses. By mining a large array of ac^4^C epitranscriptome datasets with deep learning-based pipelines, the OpenAc4C offers a comprehensive collection of ac^4^C-related resources including: (i) an ac^4^C database holding the most complete collection of 536,745 ac^4^C sites spanning a total of 33 species. It includes four distinct ac^4^C profiling techniques based on NGS platform and represents the first attempt to collect 88,936 putative ac^4^C sites derived from ONT-based studies; (ii) a genetic variants database consisting of 536,986 ac^4^C-affecting variants predicted by deep-learning pipelines in seven species, including 4,766 human disease-associated TagSNPs that may function at the epitranscriptome layer through ac^4^C disturbance; (iii) a data annotation feature that systemically integrates basic annotation with functional downstream regulation such as RBP-binding, microRNA interaction and splicing events; (iv) and a web-based analysis platform allowing interactive processing of the database collections and user-uploaded genetic variant datasets. OpenAc4C also provides a comprehensive graphical visualization and web interface for visually displaying the collected data. Together, we expect that OpenAc4C will serve as a valuable integrated resource to facilitate studies of ac^4^C modification (**Figure 1**). It is freely accessible at: www.rnamd.org/ac4cportal.

**Figure 1.**
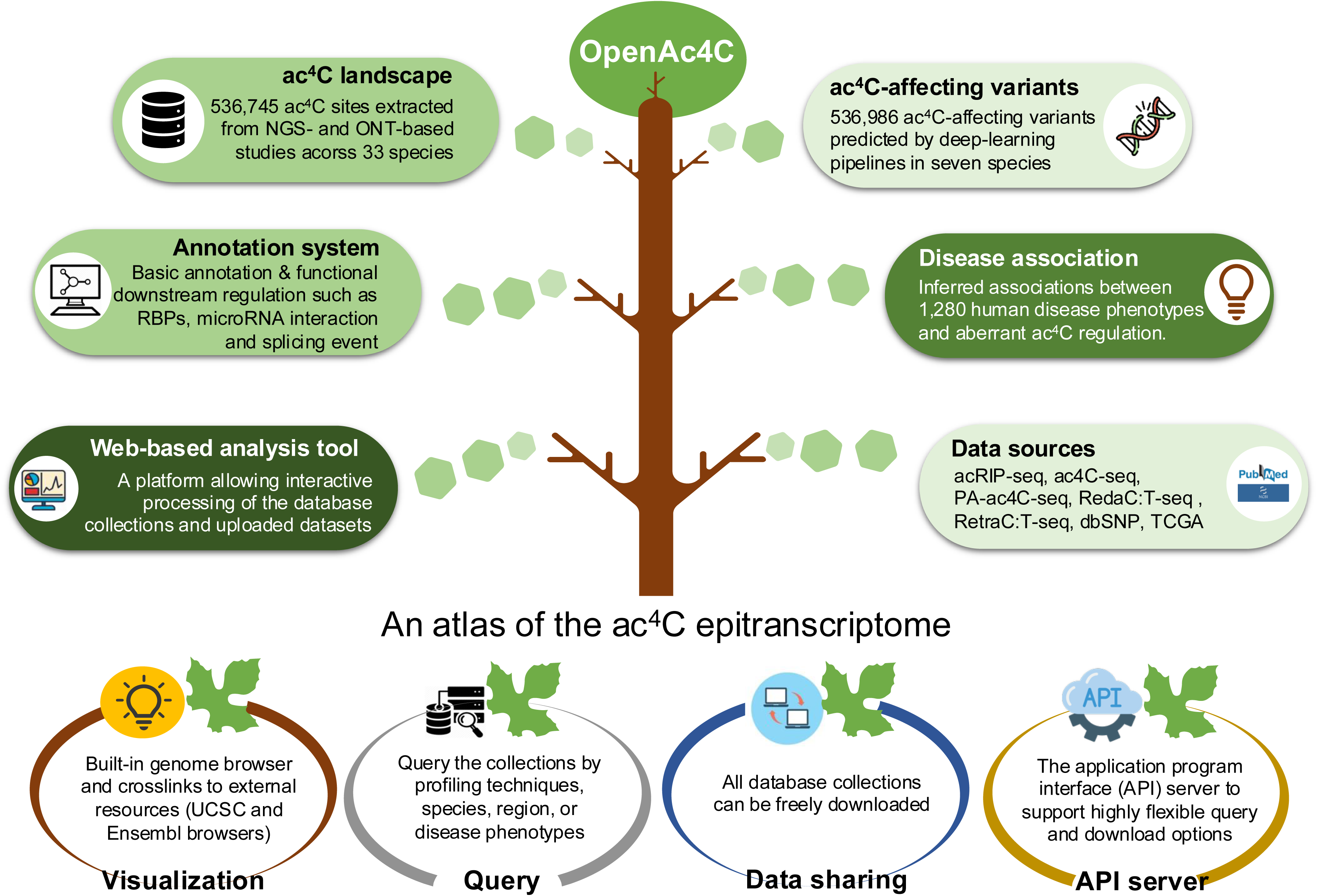
The overall design of the OpenAc4C. The OpenAc4C platform holds ∼ 530,000 previously reported ac^4^C sites identified by four distinct NGS-based ac^4^C profiling techniques, also features the first collection of putative ac^4^C sites from direct RNA sequencing samples facilitated by a deep learning model. The ac^4^C-associated variants (involving addition or removal of an ac^4^C site) were identified in seven species, among which potential associations in ∼ 1300 disease phenotypes were revealed by pathogenic ac^4^C-SNPs. In addition, two deep learning-powered web tools were presented to support analyses of the database collections and uploaded new datasets. An integrated web platform offers comprehensive query and functional exploration of all collected ac^4^C data is freely accessible at: www.rnamd.org/ac4cportal.

## Data collection and processing

### Collection of NGS-based ac^4^C modification sites

In this study, we collected and integrated hundreds of ac^4^C profiling datasets based on NGS platforms using four different sequencing techniques (**Figure S1**). A total of 226 raw sequencing samples were directly downloaded and processed from the NCBI Gene Expression Omnibus (GEO) database [47] (**Supplementary Sheet S1**). These include NGS-based ac4C-RIP-seq (∼150 bp) and base-resolution NGS samples, respectively. For ac4C-RIP-seq samples, to ensure data consistency across divergent sequencing platforms, we implemented a standardized data processing framework. Specifically, the adapter sequences from the original fastq file were removed using Trim Galore (v0.6.6). Raw reads were mapped to the reference genome using STAR (v2.7.6a) with default parameters [48], followed by peak-calling using exomePeak2 with a unified cut-off of *P*-value < 1e-10 [49]. Crucially, for datasets containing biological replicates, we performed joint peak-calling to model biological variation across samples, while incorporating GC content correction to mitigate technical biases and enhance statistical consistency. To harmonize data across studies, all identified peaks were mapped to the latest genome assemblies (e.g., hg38 for human and mm10 for mouse) using the UCSC LiftOver tool. Finally, the enriched results from all samples were merged. In addition, the genome coordinates of high resolution ac^4^C sites, including base-nucleotide mapping, were extracted from the relating GSE or corresponding supplementary files of PA-ac4C-seq [23], ac4C-seq [24, 27], and RedaC:T-seq [25] studies, respectively. Additionally, a limited number of ac^4^C acetylation sites from other public repositories were integrated into our database with labeled sources [35, 36].

### The landscape of ac^4^C acetylation unveiled by the ONT-platform

42 FAST5 and 65 FASTQ files derived from 28 independent studies were collected from the NCBI GEO database (**Supplementary Sheet S2**). First, we leveraged the ELIGOS pipeline combined with a deep learning model for a large-scale prediction of modified cytosines [50]. Initially, ELIGOS was employed to filter out chemically modified (non-canonical) cytosine bases from the direct RNA sequencing samples, based on the base-calling errors introduced by non-canonical ribonucleotide bases compared with normal ones. Specifically, raw FAST5 files were base-called by Guppy and then aligned to the reference genome using Minimap2[51]. The base call error features from the aligned SAM files were extracted, identifying sites with significantly higher errors as candidate modified cytosines. As ELIGOS reports only modified bases, without differentiating their specific modification types due to the detection of common base-calling errors [33], the modified cytosines were further assessed using our deep neural network model previously proposed [52], which was trained on experimentally-validated NGS-ac^4^C sites. Only the modified cytosines that received a prediction score > 0.5, averaged from six independent neural models (human, mouse, rat, yeast, rice, and *Arabidopsis*), and were also within the top 1% (upper bound of the *P*-value < 0.01) of all modified cytosines, were retained as putative ONT-based ac^4^C sites. In addition, we integrated DirectRM [53], a state-of-the-art tool capable of decoding RNA modifications directly from raw nanopore ionic currents. For the subset of samples with raw signal files (POD5 or FAST5 format), DirectRM was utilized to identify base-resolution ac^4^C sites by analyzing ionic signal anomalies. Please refer to **Supplementary Methods** for details.

### Decoding the epitranscriptomic effect of genetic variants on ac^4^C disturbance

To explore the potential impact of genetic variants on ac^4^C disturbance, we gathered a collection of 85,920,859 genetic variants across seven species (human, rat, mouse, zebrafish, rice, yeast, *Arabidopsis*), from which the comprehensive map of functional variants driving ac^4^C dysregulation was revealed. We considered both the germline and cancer somatic variants to perform the analysis, extracting from various resources including dbSNP (v151) [54], Ensembl 2022 (Ensembl release 106) [55], 1000 Genomes (Phase3 Mitochondrial Chromosome Variants set), and the Cancer Genome Atlas (TCGA) (release version v35) [56]. The detailed information of variant datasets is listed in **Supplementary Sheet S3**.

To systematically identify the genetic drivers within the epitranscriptome layer circuitry, we uncovered maps of functional variants by examining their ability to drive ac^4^C dysregulation, based on the prediction results for reference and mutated sequences using our previously proposed deep-learning approach [52]. Inspired by RMVar [42] and RMDisease [43], ac4C-affecting variants were categorized into two major classes based on their impact on ac^4^C status. ac^4^C-loss variants are mutations that result in the removal of modified residues, either by directly destroying an experimentally validated ac^4^C sites at base-resolution level (ac4C-seq, RedaC:T-seq, and RetraC:T-seq; high confidence level), or by altering a nucleotide within the identified ac^4^C-enriched regions (acRIP-seq; medium confidence level), significantly decreasing the ac^4^C probability as predicted by a deep neural model. In addition, we performed a transcriptome-wide prediction for all canonical cytosines, evaluating their association level (AL) based on ac^4^C probabilities in wild-type (<$>P_{WT}<$>) and mutated (<$>P_{SNP}<$>) sequences from Equation (1). The significantly increase (ac^4^C-gain variants) or decrease (ac^4^C-loss variants) in predicted ac^4^C probabilities was calculated and classified into a low confidence level, representing a transcriptome-wide prediction. The AL ranges from 0 to 1, indicating the increasing impact (association) of a mutation on ac^4^C acetylation status. Consequently, only the ac^4^C-associated variants within the absolute ranking of top 1% ALs of all mutations (upper bound of the *P*-value < 0.01) were retained in the database collection.

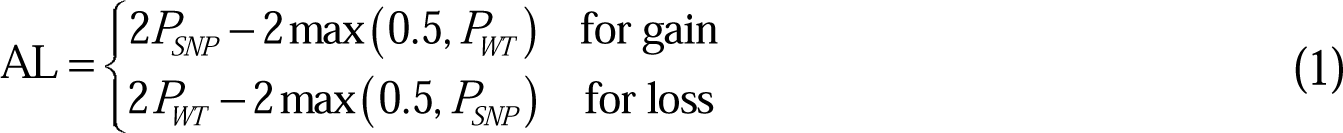

### Data annotation and visualization

A data annotation and visualization system were applied to the database collections, featuring six different annotation modules to support comprehensive exploration of the collected ac^4^C modification sites. (i) the data sources module clearly presents the driving source of the ac^4^C sites extracted from experimentally validated studies, including PubMed ID, profiling techniques, GEO datasets (GEO number), cell lines/tissues, and experimental treatment. (ii) the gene annotation module annotates the genomic coordinates of ac4C sites, including genome assembly and genomic coordinates (chromosome, position, and strand). It also provides gene information such as gene symbol, gene type, transcript region, Ensembl gene ID, and the extracted ac^4^C-containing sequences. (iii) the data reproducibility module to present both sample reproducibility and technique reproducibility for each collected ac^4^C site. (iv) the data visualization module offers a built-in Jbrowse genome browser [57], along with links to external resources such as the UCSC genome browser [58] and Ensembl coordinate mappings [55], for interactive exploration of genome coordinated-based records. Additionally, graphical visualization of the predicted RNA secondary structure was generated by RNAfold [59]. (v) The post-transcriptional regulation module integrates data from POSTAR2 (RBP binding regions) [60], miRanda and starBase2 (miRNA-RNA interactions) [61, 62], and UCSC browser annotations (GT-AG splicing sites) [58]. In addition to the functional modules mentioned above, (vi) the mutation annotation module presents the collected ac4C-associated variants, offering various annotations such as TCGA projects, TCGA barcodes [56], RS ID [54], mutation type (nonsynonymous or synonymous variants) [63], deleterious level [63], predicted ac4C-affecting function (ac4C-gain or ac4C-loss function), and genome visualization.

### Exploring the association between ac^4^C disturbance and disease pathogenesis

The potential involvement of ac^4^C in various diseases was evaluated through a systematic assessment of disease-relevant genetic mutations (disease-TagSNPs) potentially associated with ac^4^C dysregulation. We integrated data from multiple comprehensive resources, including the NCI Genomic Data Commons (GDC) Data Portal [64], GWAS catalog [65], ClinVar [66], and Johnson and O’Donnel’s database [67], to identify numerous disease-associated variants. Furthermore, the deleteriousness of these disease-TagSNPs was analyzed using five independent scoring systems [68–71]. To ensure high-confidence mapping, we required an exact genomic coordinate match between a known disease TagSNP and a predicted ac^4^C-affecting variant (all three confidence level). These prioritized variants were then used to unveil potential mechanisms of disease phenotypes that may be linked to the perturbation of the ac^4^C epitranscriptome.

### Implementation of OpenAc4C database and web interfaces

All the processed datasets, enriched with annotations, are presented on OpenAc4C, an integrated online platform constructed with Hyper Text Markup Language (HTML), Cascading Style Sheets (CSS), Javascript, and Hypertext Preprocessor (PHP). External packages and integrated frameworks have been applied to present metadata and enhance the website’s layout, including MySQL, Echarts, Bootstrap, and DataTable. Additionally, R and Python scripts were utilized to develop built-in web servers, enabling diverse interactive analyses to be performed on the database collections.

## Database content and usage

### An atlas of ac^4^C modification sites across 33 species

To comprehensively unveil the global map of the ac^4^C epitranscriptome, the OpenAc4C database contains a total of 536,745 ac^4^C sites. The data in OpenAc4C consists of four distinct NGS-based experimental techniques (acRIP-seq, PA-ac4C-seq, ac4C-seq, and RedaC:T-seq) alongside deep-learning-powered prediction derived from Oxford Nanopore (ONT) direct RNA sequencing (see **Table 1**). The NGS-based ac^4^C sites were derived from 14 species spanning archaea, virus, and eukarya, including human (207,917 sites), mouse (101,544 sites), rat (8,110 sites) , *Arabidopsis* (38,085 sites), zebrafish (4,940 sites), yeast (120 sites), rice (74,449 sites), nematode (9,011 sites), *Thermococcus kodakarensis* (404 sites), *Sulfolobus solfataricus* (45 sites), *Pyrococcus furiosus* (232 sites), *Thermococcus* sp.AM4 (266 sites), and Kaposi’s sarcoma-associated herpesvirus (KSHV, 274 sites). Among them, base-level profiling techniques were applied to eight species, covering human (2,011 sites identified by ac4C-seq and 13,481 sites by RedaC:T-seq, 1bp resolution), yeast (120 sites, ac4C-seq, 1bp resolution), rice (67,655 sites, ac4C-seq, 1bp resolution), *Arabidopsis* (34,440 sites, ac4C-seq, 1bp resolution), and the four types of archaea (a total of 947 sites, ac4C-seq, 1bp resolution). The ac^4^C landscape in the remaining species was profiled by acRIP-seq, targeting ac^4^C-enriched region with a resolution of approximately 150 bp. For ac^4^C sites identified in direct RNA sequencing samples, we collected a total of 88,936 ac^4^C modified cytosines that passed our selection criteria (please refer to **Supplementary Methods**). Each of these modified cytosines was assigned an ac^4^C score (a probability value), representing putative ONT-based ac^4^C sites across 20 species. In summary, OpenAc4C uniquely collects ac^4^C sites from 24 species for the first time and enables the timely sharing of ac^4^C data derived from direct RNA sequencing studies. A comparison between OpenAc4C and other epitranscriptome databases is detailed in **Table 2**.

**Table 1.** Collection of ac^4^C sites in OpenAc4C.

| Species | Experimentally-derived<br>NGS techniques |  | ONT-derived and<br>deep-learning prediction | Total |
| --- | --- | --- | --- | --- |
|  | 1bp | 10~150bp | 1bp |  |
| Human | 15,487 | 192,430 | 45,966 | 253,883 |
| Mouse | / | 101,544 | 4,970 | 106,514 |
| Rat | 24 | 8,086 | / | 8,110 |
| Zebrafish | / | 4,940 | / | 4,940 |
| Pig | / | / | 397 | 397 |
| Yeast | 120 | / | 32 | 152 |
| Rice | 67,655 | 6,794 | / | 74,449 |
| *Arabidopsis* | 34,440 | 3,645 | 2,893 | 40,978 |
| Worm | / | 9,011 | / | 9,011 |
| Archaea (four species) | 947 | / | / | 947 |
| KSHV | / | 274 | / | 274 |
| 19 other species | 50 | 2,362 | 34,678 | 37,090 |
| Total | 118,723 | 329,086 | 88,936 | 536,745 |

**Note:** In humans, NGS-based base-resolution ac<sup>4</sup>C sites were identified using RedaC:T-seq and ac4C-seq, ONT-based ac<sup>4</sup>C sites were predicted by ELIGOS and DirectRM software (please refer to METHODS for details). Only ac4C-seq was employed to detect base-resolution ac<sup>4</sup>C sites in other species. The overlapped ac<sup>4</sup>C sites between different samples and sequencing techniques can be checked in the ‘Data Reproducibility’ module in the OpenAc4C database.

**Table 2.** Comparisons between OpenAc4C and existing epitranscriptome databases.

|  | OpenAc4C<br>v1.0 | RMBase<br>v3.0 | RMDisease<br>v2.0 | RMVar | DirectRM<br>DB |
| --- | --- | --- | --- | --- | --- |
| Database collection (ac<sup>4</sup>C sites) | 536,745 | 2,077 | 9,365 | / | 3 |
| Number of covered species | 33 | 12 | 1 | / | 1 |
| Database collection (ac<sup>4</sup>C-SNP) | 536,986 | / | 24,049 | / | / |
| Number of covered species | 7 | / | 1 | / | / |
| Pathogenic ac<sup>4</sup>C-SNP | 4,766 | / | 874 | / | / |
| Related disease phenotypes | 1,280 | / | 299 | / | / |
| Web-based analysis tool for<br>identification ac<sup>4</sup>C site | Yes<br>(13 species) | / | / | / | / |
| Web-based analysis tool for<br>identification ac<sup>4</sup>C-SNP | Yes<br>(6 species) | / | Yes<br>(1 species) | / | / |

**Note:** The complete incorporation of currently available ac<sup>4</sup>C profiling datasets makes our proposed database the most comprehensive ac<sup>4</sup>C knowledgebase to date. Previous databases, such as RMBase, only cover a very limited number of base-resolution ac<sup>4</sup>C sites. The number of covered species has increased from 12 in RMBase to 33 in OpenAc4C. Additionally, the incorporation of ONT-based datasets serves as a valuable supplement to NGS collections.

The integration of extensive ac^4^C datasets allows us to independently assess data reproducibility across different sequencing samples. First, we examined the overall data reproducibility between ac^4^C sites generated by base-resolution techniques and acRIP-seq. In human, we found that 5,897 out of 13,481 (43.74%) base-resolution ac^4^C sites generated by RedaC:T-seq can be independent detected by the ac^4^C-enriched peaks using acRIP-seq (**Figure S2A**). In addition to RedaC:T-seq, this number is 1,220 out of 2,011 (60.67%) for ac4C-seq (**Figure S2B**). We then calculated the probability of these co-occurrence happening completely by chance to assess the statistical significance of our findings. Specifically, we randomly selected 15,487 pseudo ac^4^C sites from the cytosines on the same transcripts of all base-resolution ac^4^C sites (RedaC:T-seq and ac4C-seq). These pseudo ac^4^C residues were then mapped to the same ac^4^C-enriched peaks to count overlapped sites. This process was repeated 1,000 times. The analysis result indicated that the number of base-resolution ac^4^C sites independently detected by acRIP-seq (7,115 out of 15,487; 45.94%) is significantly greater than what would be expected by chance (481 out of 15,487; 3.1%, **Figure S2C**). Besides the different sequencing approaches generated on NGS-platform, we then examined the overlapped sites between ONT-derived prediction and NGS-based experimental collection. The analysis was performed on three species (human, mouse, and *Arabidopsis*) that encompass a relatively considerable number of ac^4^C sites generated from both platforms. For human, we found that 18,167 out of 45,966 (39.52%) human ONT-derived ac^4^C sites can be independently detected using at least one NGS-based technique (**Figure S3A**). In addition to human, this number is 1,795 out of 4,970 (36.12%, **Figure S3B**) for mouse and 185 out of 2,893 (6.39%, **Figure S3C**) for *Arabidopsis*, respectively. A permutation test confirmed that the observed human overlap is significantly greater than what would be expected by chance (**Figure S3D-S3F**), suggesting that the ONT-based ac^4^C sites collected in our database serve as a valuable supplement to NGS collections.

### Associations between ac^4^C dysregulation and diseases revealed by ac^4^C-affecting variants

Mutations linked to aberrant epitranscriptomic machinery are involved in various disease and cancer [72, 73]. OpenAc4C provides a comprehensive map for studying the genetic drivers that may operate through aberrant ac^4^C machinery. A total of 536,986 ac^4^C-associated genetic variants (which add or remove ac^4^C acetylation sites) were collected across seven species (**Table 3**), covering human (257,553), rat (8,113), mouse (177,368), rice (19,588), *Arabidopsis* (40,246), zebrafish (34,107), and pig (11). The associations between disease pathogenesis and ac^4^C regulation were revealed by mapping human ac^4^C-associated variants to disease TagSNPs with pathogenic records. We found 4,766 disease-relevant potentially ac^4^C-associated variants that localized on 1,851 genes and linked to 1,280 known disease phenotypes. In addition, the implications of ac^4^C disturbance in cancer were revealed by 165,081 somatic ac^4^C-associated variants, deriving from 33 types of TCGA cancer projects.

**Table 3.** ac^4^C-associated variants and their disease relevance.

| Species | Confidence Level | ac<sup>4</sup>C-associated variants |  |  | ClinVar |  |  | GWAS |  |  |
| --- | --- | --- | --- | --- | --- | --- | --- | --- | --- | --- |
|  |  | Loss | Gain | Total | SNP | Disease | Gene | SNP | Disease | Gene |
| Human (Germline SNP) | High | 598 | / | 598 | 52 | 42 | 52 | 4 | 4 | 4 |
|  | Medium | 16,787 | / | 16,787 | 734 | 335 | 339 | 89 | 58 | 73 |
|  | Low | 37,227 | 37,923 | 75,150 | 2,413 | 822 | 981 | 533 | 233 | 486 |
| Human (Somatic SNP) | High | 3,184 | / | 3,184 | 30 | 22 | 27 | / | / | / |
|  | Medium | 14,485 | / | 14,485 | 88 | 54 | 65 | / | / | / |
|  | Low | 89,454 | 57,895 | 147,349 | 891 | 481 | 455 | / | / | / |
| Mouse | High | 19 | / | 19 | / | / | / | / | / | / |
|  | Medium | 2,564 | / | 2,564 | / | / | / | / | / | / |
|  | Low | 90,760 | 84,025 | 174,785 | / | / | / | / | / | / |
| Rat | Medium | 265 | - | 265 | / | / | / | / | / | / |
|  | Low | 3,913 | 3,935 | 7,848 | / | / | / | / | / | / |
| Pig | High | 11 | / | 11 | / | / | / | / | / | / |
| Zebrafish | Medium | 2,364 | / | 2,364 | / | / | / | / | / | / |
|  | Low | 15,775 | 15,968 | 31,743 | / | / | / | / | / | / |
| Rice | High | 955 | / | 955 | / | / | / | / | / | / |
|  | Medium | 4,283 | / | 4,283 | / | / | / | / | / | / |
|  | Low | 7,132 | 7,218 | 14,350 | / | / | / | / | / | / |
| Arabidopsis | High | 2,032 | / | 2,032 | / | / | / | / | / | / |
|  | Medium | 9,332 | / | 9,332 | / | / | / | / | / | / |
|  | Low | 14,923 | 13,959 | 28,882 | / | / | / | / | / | / |

Note: High confidence level: genetic variants directly destroying base-resolution ac<sup>4</sup>C modified nucleotides. Medium confidence level: genetic variants altering a nucleotide within the identified ac<sup>4</sup>C-enriched regions. Low confidence level: the transcriptome-wide prediction for reference- and mutated-sequence (altered by a mutation) around cytosine, including all predicted ac<sup>4</sup>C-gain variants.

We then calculated the predominant disease phenotypes and TCGA cancer types with a significant correlation to aberrant ac^4^C status (**Table 4**). Among them, 116 pathogenic ac^4^C-associated variants (2.43%) were related to hereditary cancer-predisposing syndrome (ClinVar study, Accession: RCV000494198.1). These were followed by 57 variants (1.20%) associated with nonsyndromic hearing loss (ClinVar study, Accession: RCV000322137.1) and 47 variants (0.99%) linked to thoracic aortic aneurysm and aortic dissection (ClinVar study, Accession: RCV000249596.1). Additionally, we found that 279 ac^4^C-associated variants were also recorded as cancer-somatic variants in the Cholangiocarcinoma (TCGA-CHOL) project, representing 8.40% of the total TCGA-CHOL somatic mutations and ranking as the top enriched TCGA cancer type, followed by Esophageal Carcinoma (TCGA-ESCA) and Uterine Corpus Endometrial Carcinoma (TCGA-UCEC) as the second and third most enriched TCGA cancer types, respectively.

**Table 4.** Disease phenotypes and TCGA cancer projects that most enriched with aberrant ac^4^C status.

| ClinVar |  |  |  |  |  | TCGA |  |
| --- | --- | --- | --- | --- | --- | --- | --- |
| Name | # | Database | Study accession | Clinical significance | Identifier | Project name | Ratio* |
| Hereditary cancer-predisposing syndrome | 116<br>(2.43%) | ClinVar | RCV000494198.1 | Pathogenic | MedGen:<br>C0027672 | Cholangiocarcinoma<br>(TCGA-CHOL) | 8.40% |
| Nonsyndromic hearing loss | 57<br>(1.20%) | ClinVar | RCV000322137.1 | Likely<br>Pathogenic | MedGen:<br>CN239439 | Esophageal<br>Carcinoma<br>(TCGA-ESCA) | 8.29% |
| Thoracic aortic aneurysm and aortic dissection | 47<br>(0.99%) | ClinVar | RCV000249596.1 | Pathogenic | MedGen:<br>CN118826 | Uterine Corpus<br>Endometrial<br>Carcinoma<br>(TCGA-UCEC) | 7.99% |

Ratio\*: ac<sup>4</sup>C-SNPs/total somatic SNPs extracted from the corresponding TCGA project.

### Enhanced web interface and application

OpenAc4C features an informative and user-friendly web interface with comprehensive query options, customized searches, graphic visualizations, and batch downloading of all database collections, offering an integrated knowledgebase to facilitate research in ac^4^C community.

#### Query ac^4^C sites with rich annotations

The collected ac^4^C sites in OpenAc4C are classified by profiling techniques. The ‘NGS High Resolution’ module contains the base-resolution ac^4^C collection profiled by NGS techniques. The ‘NGS acRIP-seq’ module holds the ac^4^C-enriched regions detected by acRIP-seq, and the ‘Direct RNA Sequencing’ module to unveil the landscape of ac^4^C acetylation generated by direct RNA sequencing samples. By entering the corresponding repository, users are presented with three intuitive pie charts that provide a general summary of the collected data, categorized by species, gene region, and gene type. This is accompanied by various functional filters on the left panel, allowing users to further extract data of interest. The database table retrieves the ac^4^C datasets that satisfy all customized filter options, and detailed information can be accessed by clicking a specific site ID, including genomic coordinates, data source, profiled condition, and potential involvement in post-transcriptional regulation. The ‘Data Reproducibility’ module indicates whether this ac^4^C site can be independently detected by different sequencing samples or techniques. The ‘Match ID’ links to the detailed pages of overlapped records (**Figure 2**).

**Figure 2.**
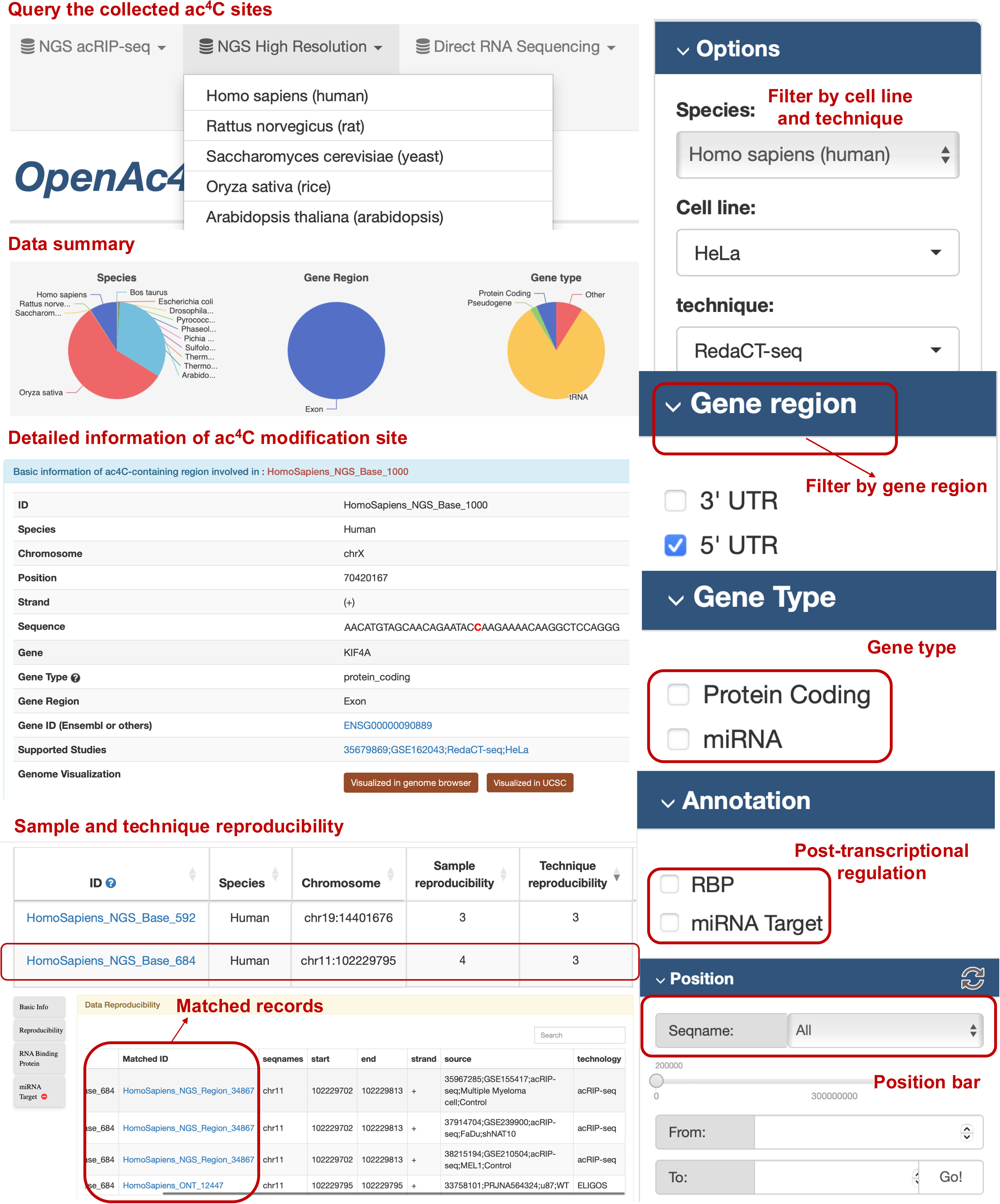
ac^4^C landscape of OpenAc4C. The collected ac^4^C sites were classified by their profiling techniques. Users can easily check the statistical distribution of collected data summarized by pie charts. Various filter options are provided to filter data of interest. Detailed information for each ac^4^C site can be accessed by clicking on a specific site ID, including genome coordinate, sequence, gene annotation, deriving source (experiment), and data reproducibility.

#### Explore disease phenotypes associated with ac^4^C disturbance

The ac^4^C-associated variants and disease associations were collected in ‘ac4CDiseaseDB’ module and can be explored in two ways (**Figure 3**). (i) the collected ac^4^C-associated variants can be accessed by clicking the ‘ac4CDiseaseDB’ button on the top navigation bar. The main data table presents the basic information of collected ac^4^C-SNPs, such as species, genome coordinates, modification status, and confidence level, and changes accordingly based on the customized filter options. The disease associations can be obtained by clicking ‘GWAS’ or ‘ClinVar’ buttons from the filter columns, followed by clicking a specific RM ID to view detailed information such as disease phenotype, ClinVar/GWAS accession number, reference/mutated sequence, potential involvement in post-transcriptional regulation, and their predicted impact on ac^4^C disturbance. (ii) additionally, to facilitate querying a specific disease phenotype, keywords can be directly entered in the homepage search box using ‘Disease’ searching mode. The pages will return a full list of relevant results linking all database collections to the searched keywords (disease phenotype). To support graphic visualization, OpenAc4C also provides a built-in genome browser and crosslinks to external resources (UCSC and Ensembl browsers) for interactive navigation of ac^4^C-SNPs and their affecting genes (regions).

**Figure 3.**
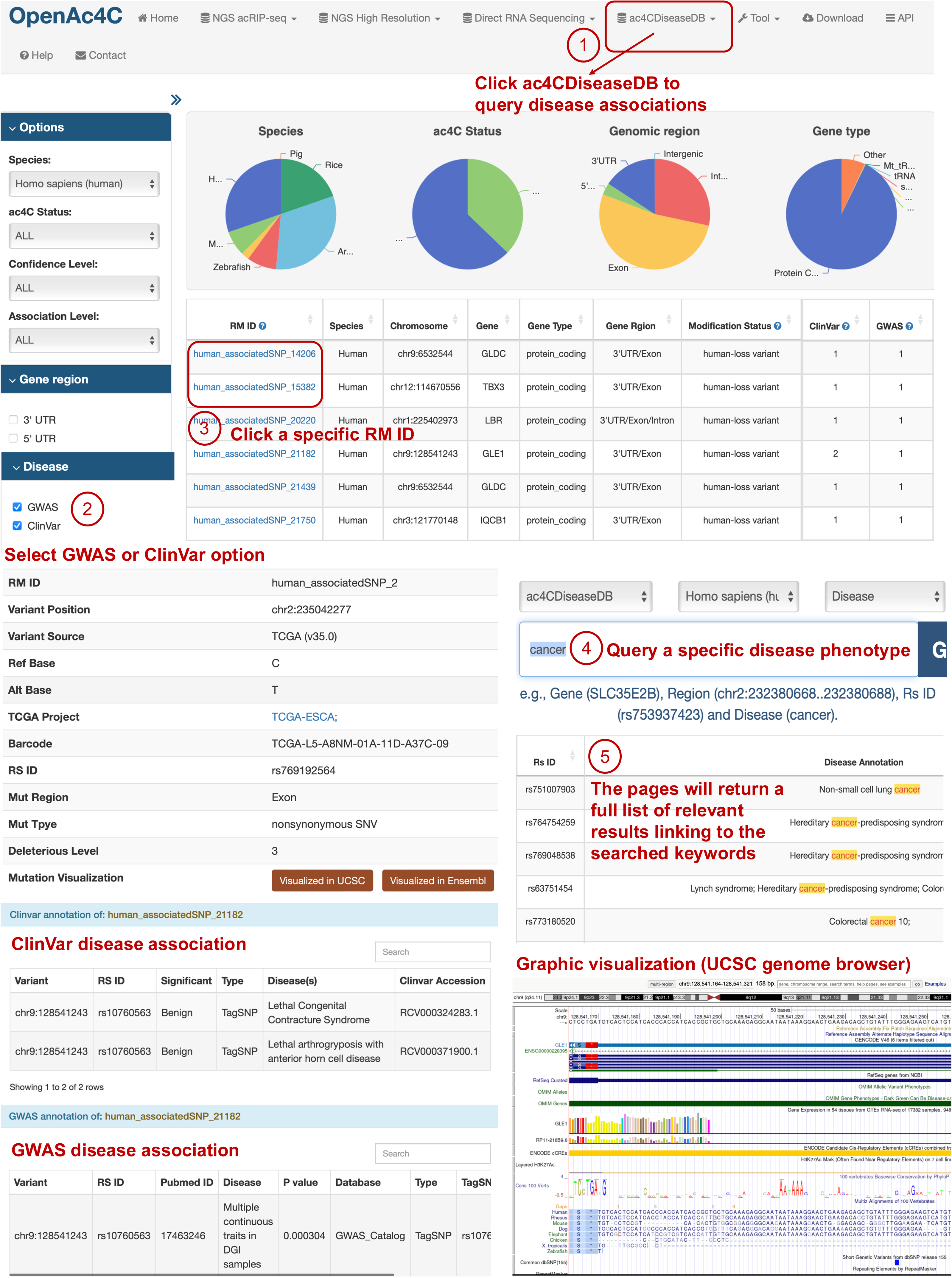
The disease associations revealed by ac^4^C-associated variants. The disease associations revealed by ac^4^C-associated variants can be explored by users through the ‘ac4CdiseaseDB’ accessible via the top navigation bar. By activating the ‘ClinVar’ or ‘GWAS’ filter options, the search results will specifically highlight ac^4^C sites linked to diseases. Users can click on an individual RM ID to access detailed information about the associated ac^4^C-SNP and its correlated disease. Beyond general queries, the platform supports customized searches for investigating specific disease phenotypes, encompassing all related data compiled. The web interface is further enhanced by various graphical visualizations, including Ensembl and UCSC genome browsers, allowing for interactive exploration of genomic regions of interest.

#### Web-based analysis modules

Two data analysis tools have been developed to enable users to perform interactive analyses on the database collections or user-uploaded datasets. (i) A deep learning-powered prediction module supports *in-silico* identification of putative ac^4^C sites from RNA sequence inputs (in standard FASTA format) across 13 species. This module is built on our previously developed multi-instance learning framework [52] (**Supplementary Method**). Instructions on preparing input data, selecting prediction parameters, and interpreting results are provided with clear guidance (**Figure 4A-B**). If an incorrect input format is detected, the server will automatically terminate the process and alert the user (**Figure 4C**). (ii) An enhanced SNP analysis module allows users to upload a set of genetic variants (in standard VCF format) for interactive analysis with experimentally-derived ac^4^C sites and transcriptome-wide cytosines. The deep learning model evaluates the ac^4^C probability on both reference and mutated sequences. Newly identified ac^4^C-affecting SNPs are labeled with their predicted ac^4^C function (ac^4^C-gain or ac^4^C-loss), potential affected region, association level (AL), and confidence level (**Figure 4D**). All results from our web-based analysis tools can be downloaded as a batch file.

**Figure 4.**
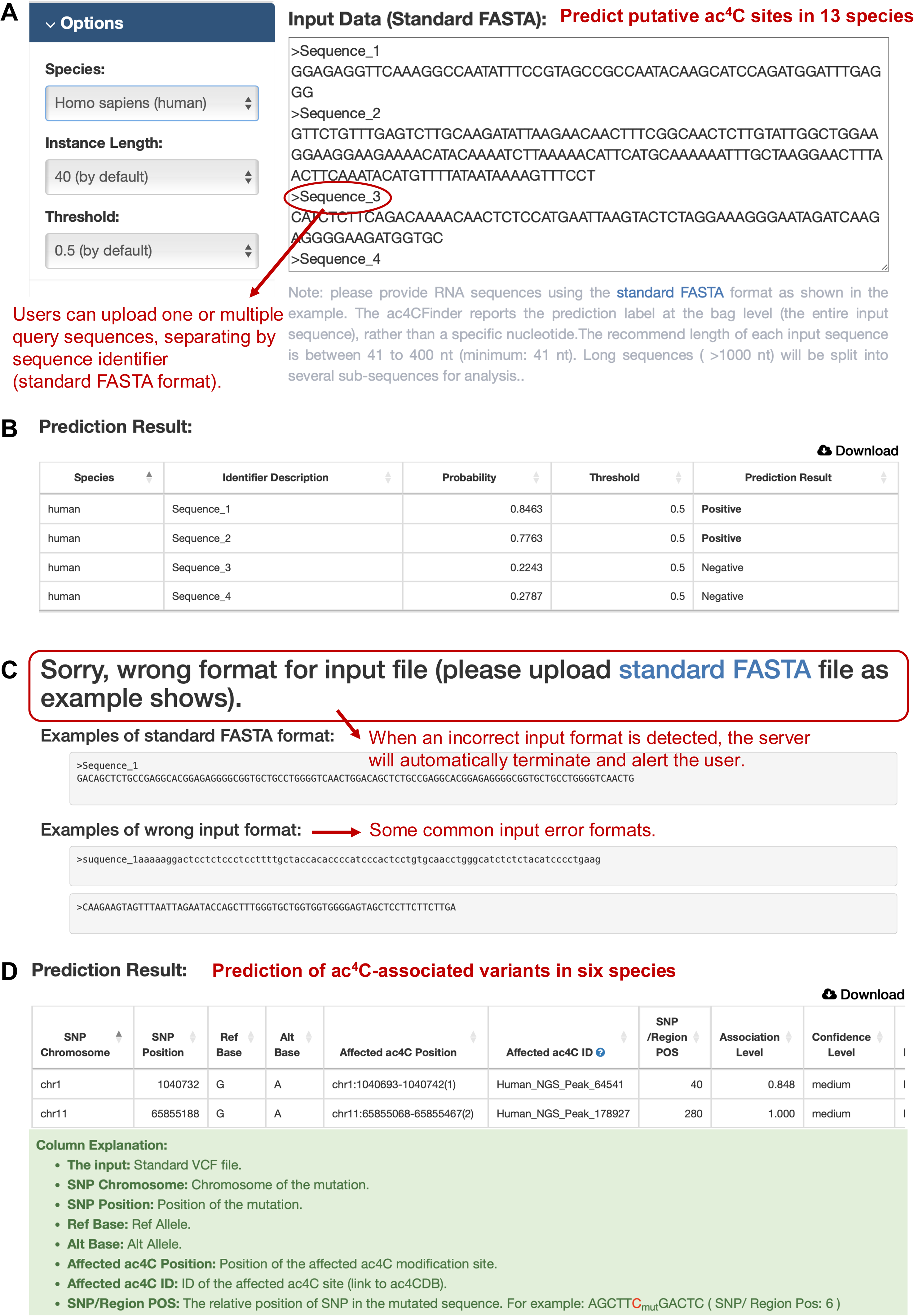
Web-based analysis tools. **A**. Input data format and parameters required for predicting potential ac^4^C sites across 13 species. **B**. The server generates an output report detailing the predicted probability that each input sequence undergoes ac^4^C modification. **C**. Users will receive an error notification prompting them to prepare the input in the correct format. **D**. The prediction output for ac4C-impacting variants includes a column with explanations.

#### Data downloading and sharing

All database collections can be freely downloaded categorized by profiling techniques, data resolution, detected platform, and covered species. Multiple datasets can be simultaneously selected for batch downloading in Comma Separated Values (CSV) format. OpenAc4C also provides an application program interface (API) server to support highly flexible query and download options. Please refer to the ‘API’ page for detailed descriptions and demo examples on how to access the API server.

## Discussion

Recent progress in ac^4^C acetylation has broadened our understanding of its substantial regulatory effects on a wide range of cellular functions and its close association with various diseases, including cancer. A centralized bioinformatics resource with a user-friendly gateway is urgently needed to gather, annotate and share the rapidly accumulating ac^4^C sequencing data. To this end, we present OpenAc4C, an integrated knowledge base that encompasses the most extensive collection of ac^4^C-related data to date (**Table 2**). Compared to other related epitranscriptome databases, OpenAc4C features a comprehensive repository of 536,745 previously reported ac^4^C modification sites across 33 species. These include the integration of four distinct NGS-profiling techniques and the first attempt to leverage deep learning power to predict putative ac^4^C sites from direct RNA sequencing samples. Beyond exploring the ac^4^C landscape, the potential impact of genetic variants on ac^4^C epitranscriptome was systemically investigated across seven species. Specifically, it revealed potential associations between 1,280 human disease phenotypes and aberrant ac^4^C regulation in 1,851 genes, identified through 4,766 pathogenic ac^4^C-associated variants. Additionally, OpenAc4C offers web-based tools for interactive analysis of database collections, allowing users to identify putative ac^4^C sites in their region of interest. It also allows for the evaluation of the associations between certain sets of genetic mutations and the collected ac^4^C sites.

It should be noted that there are still some discrepancies in detecting ac^4^C in cellular mRNA. In 2018, the acRIP-seq technique was introduced as the first method for transcriptome-wide identification of ac^4^C-enriched peaks in human mRNA [19]. This technique has been widely applied to detect mRNA ac^4^C levels across various species [20, 21], including research on human cancer [17, 74] and cellular mRNAs of KSHV transcripts [12]. In 2020, ac4C-seq was developed to quantitatively map ac^4^C sites at a single-nucleotide level [24]. It reported the detection of ac^4^C in mRNA from HEK-293T cells only under conditions where NAT10 was overexpressed. This discrepancy may be due to the variations in detection sensitivity across different profiling approaches, considering LC-MS/MS [2, 19, 27] and other transcriptome-wide profiling techniques such as RedaC:T-seq [25], RetraC:T-seq [26], and (PA)-ac4C-seq [23] have consistently confirmed the presence of ac^4^C in mRNA. We have summarized key studies related to the detection and functional exploration of mRNA ac^4^C sites across different species (**Table S1**). Nevertheless, we conducted additionally wet-lab experiments for independent validation of mRNA ac^4^C presence (**Supplementary Method**). First, we performed LC-MS/MS to measure ac^4^C levels in total RNA and oligo(dT)_25_-purified RNA from four different human cell lines: HEK293T, HT29, HCT116, and HepG2. Consistent with previous reports, ac^4^C modification was detected in all examined cell lines, with approximately 0.15% and 0.01% of the cytidines modified by ac^4^C in human total RNA and Poly(A)+ RNA, respectively (**Figure S4**). Next, to assess the consistency between different base-resolution ac^4^C profiling approaches, we employed RedaC:T-seq to identify single-base ac^4^C sites in the mRNA of the SK-Hep-1 cell line. We identified a total of 6,304 ac^4^C sites under wild-type condition and 238 sites under NAT10 knockdown condition (**Supplementary Sheet S4-S5**). We found 12 of the newly identified ac^4^C sites had been previously reported by at least one base-resolution study (**Table S2**). Furthermore, six of these 12 base-resolution sites (50%) were also identified in ac^4^C-enriched peaks profiled by ac4C-RIP-seq samples. It may be worth noting that, this limited overlap between detection methods is consistent with observations in other RNA modification type. For instance, even for the extensively studied m^6^A methylation, only approximately 10% of experimentally detected m^6^A sites is shared between two arbitrary base-resolution profiling methods [75], partially due to technical variations in experimental protocols, sequencing biases, and bioinformatic processing pipelines. Considered the relatively small number of currently reported single-base ac^4^C sites, to some degree, the consistent identification of ac^4^C sites by multiple independent base-resolution profiling methods provides a robust validation of mRNA ac^4^C presence. Our results, together, demonstrate that N4-acetylation of cytidine is a bona fide mRNA modification that can be consistently detected across various human cell lines.

Overall, an integrated ac^4^C resource will benefit the ac^4^C research community at this fast-developing stage for several reasons: (i) the ac^4^C sites identified by different profiling techniques are readily available for query and access through our database collections, offering an open platform for data sharing and reassessment. (ii) beyond human mRNA, ac^4^C has been recognized as a universally conserved modification in all domains of life. OpenAc4C provides comprehensive coverage and annotation of the ac^4^C epitranscriptome across different species and RNA types. (iii) the web-based deep-learning model offers a promising tool to facilitate the *in silico* identification and selection of potential ac^4^C sites on target mRNA regions before designing experiments.

## Conclusion

Taken together, OpenAc4C encompasses the first integrated repository of ac^4^C epitranscriptome data across 33 species. In future updates, we aim to systematically incorporate the latest sequencing datasets and emerging profiling techniques to further expand this knowledgebase. A primary focus of our future efforts will be the refinement of the ONT-based ac^4^C landscape. Our current ONT collection leverages two strategies: (i) deep learning-powered prediction based on the ELIGOS algorithm and (ii) raw signal analysis using DirectRM [53], a state-of-the-art tool capable of identifying ac^4^C directly from nanopore ionic currents. We recognize that while third-generation sequencing offers unique advantages for single-molecule and long-read profiling, the computational detection of non-m^6^A modifications from raw signals remains a significant technical challenge (Luo et al., *Nature Methods*, 2026 [76]). To ensure transparency, ONT-derived sites are currently labeled as having “limited reliability”. Nevertheless, our permutation tests (**Figure S3D-F**) demonstrate that these sites occur at a frequency significantly higher than random expectation, suggesting they provide valuable biological insights. As nanopore chemistry (e.g., RNA004) matures and more specialized signal-based algorithms become available, we will re-process our collection to enhance its precision. We remain committed to the continuous improvement of OpenAc4C and to supporting the ongoing advancement of the ac^4^C research field.

## Data availability

The raw data used to develop OpenAc4C is publicly available in the NCBI GEO database and listed in **Supplementary Sheet S1 and S2**. The datasets of germline and somatic variants derived from The Cancer Genome Atlas, dbSNP, 1000 Genome, and Ensembl 2022, are listed in **Supplementary Sheet S3**. The raw high-throughput sequencing datasets conducted for independent validation are publicly available in the BioProject database under accession number PRJNA1210697. All data collected in OpenAc4C is freely accessible at: www.rnamd.org/ac4cportal.

## CRediT author statement

**Gang Tu**: Data curation, Methodology, Software, Formal analysis, Writing – review & editing. **Yigan Zhang**: Investigation, Validation. **Xuan Wang**: Software. **Jishuai Zhang**: Investigation, Validation. **An Zhu**: Data curation. **Kunqi Chen**: Conceptualization, Writing – review & editing. **Zhixing Wu**: Data curation, Software. **Zekai Wu**: Data curation. **Yue Wang**: Writing – review & editing. **Jingxian Zhou**: Investigation, Validation. **Zhen Wei**: Data curation, Writing – review & editing. **Guifang Jia**: Writing – review & editing. **Jia Meng**: Conceptualization, Writing – review & editing. **Daniel J. Rigden**: Writing – review & editing. **Bowen Song**: Conceptualization, Data curation, Methodology, Writing – original draft, Writing – review & editing. All authors have read and approved the final manuscript.

## Supporting information

Supplementary Sheets

## Acknowledgments

National Natural Science Foundation of China [Grant No. 32500580 to B.S.]; Natural Science Foundation of Jiangsu Province [Grant No. BK20240723 to B.S.]; Scientific Research Start-up Fund for Distinguished Professor of Nanjing University of Chinese Medicine [Grant No. 013038030001 to B.S.]; Fujian Research and Training Grants for Young and Middle-aged Leaders in Healthcare (K.C.). This work is supported by the Supercomputing Platform of Xi’an Jiaotong-Liverpool University. B.S. is the academic member of the Human RNome Consortium.

## Supplementary material

**Figure S1. OpenAc4C integrates three data resolution using a rigorous, standardized pipeline.**

(i) NGS-based acRIP-seq. To ensure cross-study comparability, 226 raw samples were re-processed using a unified pipeline. (ii) NGS-based base-resolution techniques. High-resolution ac4C sites (PA-ac4C-seq, ac4C-seq, and RedaC:T-seq) were directly consolidated from author-validated records to preserve technique-specific chemical thresholds. (iii) ONT-derived and deep learning prediction. Putative ac^4^C sites from Nanopore sequencing were identified using ELIGOS and deep learning prediction. Candidates were required to meet a prediction score > 0.5 and rank within the top 1% (*P-*value < 0.01) of the modification distribution.

**Figure S2. The statistical results between acRIP-seq and base-resolution data.**

**(A)** Number of overlapped ac^4^C sites (n=5,897) between RedaC:T-seq and ac4C-RIP-seq. **(B)** Number of overlapped ac^4^C sites (n=1,220) between ac4C-seq and ac4C-RIP-seq. **(C)** Permutation test showing that the observed overlap of 7,115 unique ac^4^C sites between base-resolution and ac^4^C-enriched regions (after removing two overlapping sites between techniques) is significantly higher than the random expectation (mean = 481 sites, 1,000 iterations).

**Figure S3. The statistical results between NGS-based data and ONT-derived data.**

**(A-C)** Number of observed ac^4^C sites overlapped between ONT-derived prediction and NGS-based sequencing approaches in human (**A**), mouse (**B**), and *Arabidopsis* (**C**). (**D–F**) Permutation tests (1,000 iterations) demonstrating that the observed overlap between human ONT-derived ac^4^C sites and the consolidated NGS collection (**D**), RedaC:T-seq sites (**E**), and ac4C-seq sites (**F**) is significantly higher than the random expectation (mean values are indicated).

**Figure S4**. **LC-MS/MS analysis of four cell lines.**

**(A)** total RNA and **(B)** poly(A) RNA on HEK293T, HT29, HCT116, and HepG2, respectively. Mean ±SD, n=2 biological replicates x 2 technical replicates.

**Table S1. Techniques developed for ac^4^C mapping across various species**

**Table S2. Data reproducibility across different single-base sequencing studies**

