## Supplementary Materials for "OpenAc4C: A gateway to decode the landscape, regulation and pathogenesis of N4-acetylcytidine (ac^4^C) epitranscriptome"

**Table S1. Techniques developed for ac^4^C mapping across various species**

| Technique | Resolution | Date | mRNA presence | Species | Reference |
| --- | --- | --- | --- | --- | --- |
| FMPLA | Single-cell imaging | 2025 | Yes | Human | *Nucleic Acids Research,* [1] |
| ac4C-RIP-seq | ~150 bp | 2025 | Yes | Mouse | *Nature Immunology,* [2] |
| ac4C-RIP-seq | ~150 bp | 2025 | Yes | Human | *European Heart Journal,* [3] |
| RedaC:T-seq  ac4C-RIP-seq | 1bp  ~150 bp | 2025 | Yes | Mouse | *PNAS,* [4] |
| RetraC:T-seq | 1 bp | 2024 | Yes | Human | *RNA*, [5] |
| ac4C-RIP-seq | ~150 bp | 2024 | Yes | Human | *Nature Cell Biology*, [6] |
| ac4C-seq | 1 bp | 2024 | Yes | Rice | *Nature Plant*,  [7] |
| ac4C-seq | 1 bp | 2023 | Yes | Human | *Nucleic Acids Research*, [8] |
| ac4C-RIP-seq | ~150 bp | 2023 | Yes | Mouse | *Circulation Research*, [9] |
| ac4C-seq  ac4C-RIP-seq | 1 bp  ~150 bp | 2023 | Yes | *Arabidopsis*/Rice | *Molecular Plant*,  [10] |
| RedaC:T-seq | 1 bp | 2022 | Yes | Human | *Molecular Cell*,  [11] |
| ac4C-seq | 1 bp | 2020 | ac4C sites in mRNA detected under NAT10 overexpression condition | Human | *Nature*, [12] |
| (PA)-ac4C-seq | ~20bp | 2020 | Yes | HIV-1/Human | *Cell Host & Microbe*, [13] |
| Poly(A) RNA LC-MS/MS | / | 2018 | Yes | Human | *Cell*, [14] |
| ac4C-RIP-seq | ~150 bp | 2018 | Yes | Human | *Cell*, [14] |

**Table S2. Data reproducibility across different single-base sequencing studies**

| Chromosome | Position | Strand | Gene | RedaC:T  SK-Hep-1 | RedaC:T  2022 [11] | ac4C-seq [8] | acRIP-seq |
| --- | --- | --- | --- | --- | --- | --- | --- |
| chr1 | 17407627 | - | RCC2 | ✓ |  | ✓ |  |
| chr1 | 205717086 | - | NUCKS1 | ✓ | ✓ |  |  |
| chr3 | 51393206 | + | RBM15B | ✓ |  | ✓ | ✓ |
| chr6 | 13644583 | - | RANBP9 | ✓ | ✓ |  |  |
| chr6 | 158605274 | + | TMEM181 | ✓ | ✓ |  | ✓ |
| chr12 | 54237083 | - | CBX5 | ✓ | ✓ |  |  |
| chr12 | 56113014 | + | PA2G4 | ✓ |  | ✓ |  |
| chr12 | 120174088 | - | GCN1 | ✓ | ✓ |  | ✓ |
| chr12 | 122079222 | + | MLXIP | ✓ | ✓ |  |  |
| chr19 | 49933589 | + | ATF5 | ✓ |  | ✓ | ✓ |
| chr20 | 45326879 | - | SDC4 | ✓ | ✓ |  | ✓ |
| chr22 | 21763942 | - | MAPK1 | ✓ | ✓ |  | ✓ |

**Note**: ‘RedaC:T (SK-Hep-1)’ stands for the ac^4^C sites that independently identified in SK-Hep-1 using RedaC:T-seq, and ‘RedaC:T 2022’ stands for sites collected from its original study [11].

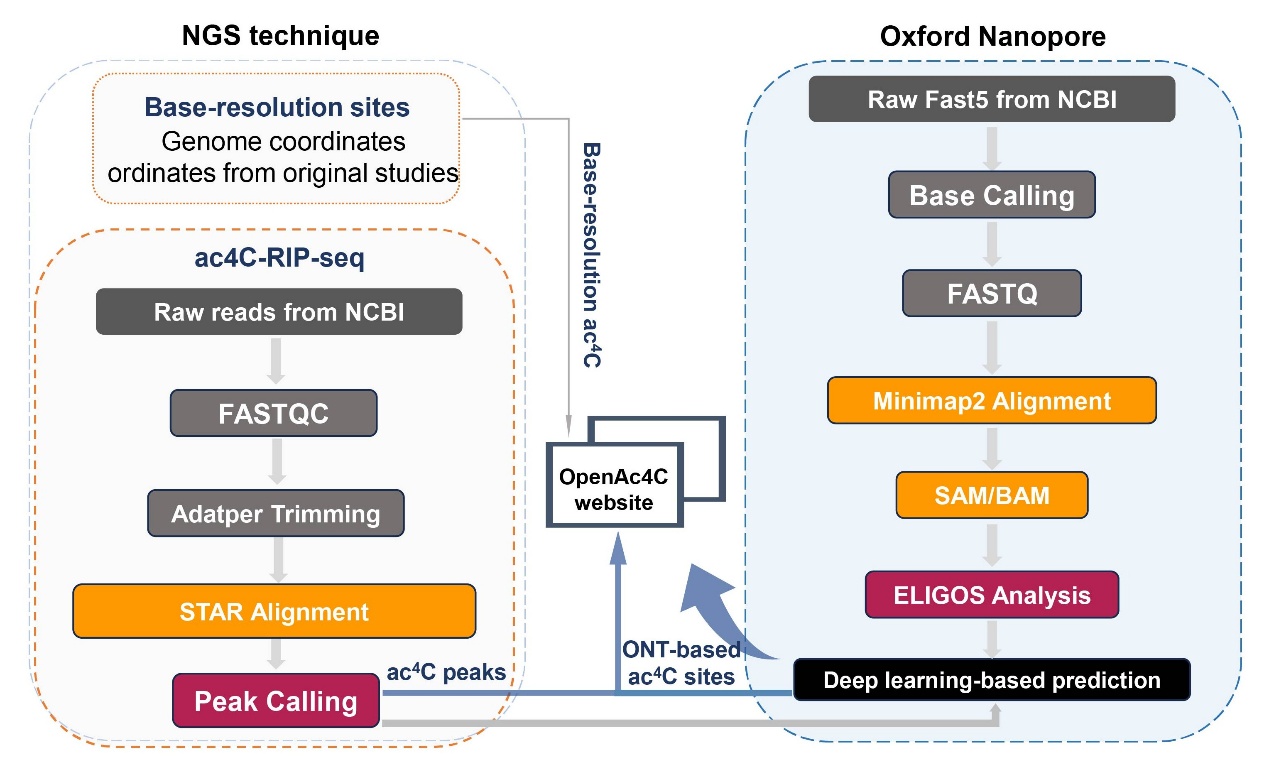

**Figure S1. OpenAc4C integrates three data resolution using a rigorous, standardized pipeline.** (i) NGS-based acRIP-seq. To ensure cross-study comparability, 226 raw samples were re-processed using a unified pipeline. (ii) NGS-based base-resolution techniques. High-resolution ac^4^C sites (PA-ac4C-seq, ac4C-seq, and RedaC:T-seq) were directly consolidated from author-validated records to preserve technique-specific chemical thresholds. (iii) ONT-derived and deep learning prediction. Putative ac^4^C sites from Nanopore sequencing were identified using ELIGOS and deep learning prediction. Candidates were required to meet a prediction score > 0.5 and rank within the top 1% (*P-*value < 0.01) of the modification distribution.

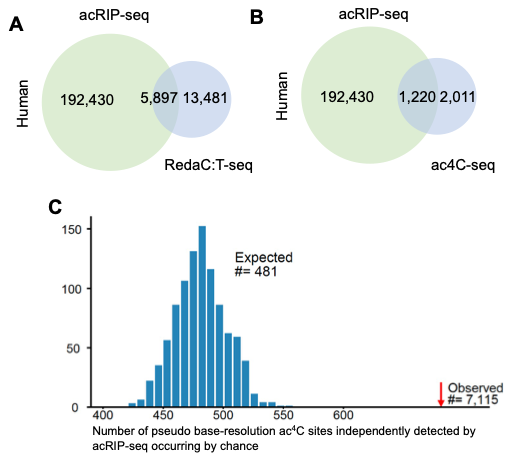
**Figure S2.** **The statistical results between acRIP-seq and base-resolution data. (A)** Number of overlapped ac^4^C sites (n=5,897) between RedaC:T-seq and ac4C-RIP-seq. **(B)** Number of overlapped ac^4^C sites (n=1,220) between ac4C-seq and ac4C-RIP-seq. **(C)** Permutation test showing that the observed overlap of 7,115 unique ac^4^C sites between base-resolution and ac^4^C-enriched regions (after removing two overlapping sites between techniques) is significantly higher than the random expectation (mean = 481 sites, 1,000 iterations).

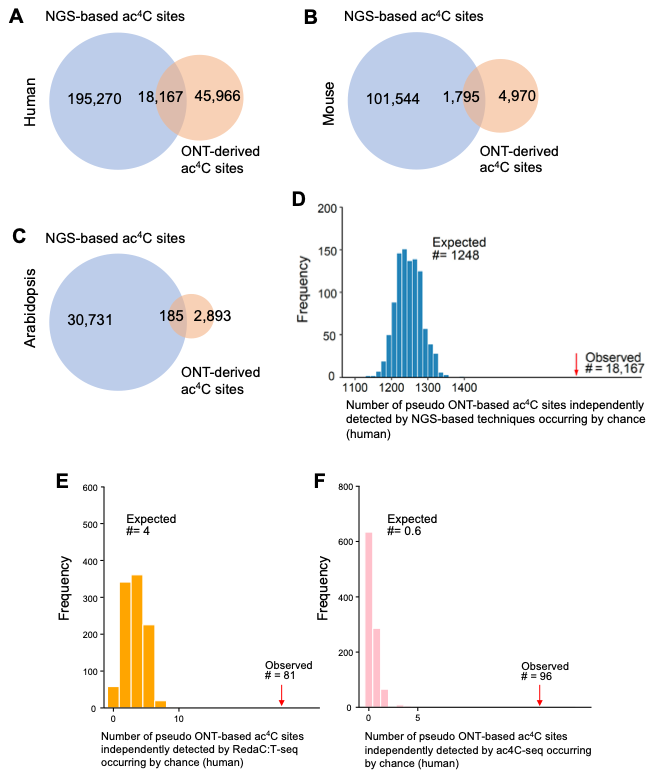

**Figure S3. The statistical results between NGS-based data and ONT-derived data. (A-C)** Number of observed ac^4^C sites overlapped between ONT-derived prediction and NGS-based sequencing approaches in human (**A**), mouse (**B**), and *Arabidopsis* (**C**). (**D–F**) Permutation tests (1,000 iterations) demonstrating that the observed overlap between human ONT-derived ac^4^C sites and the consolidated NGS collection (**D**), RedaC:T-seq sites (**E**), and ac4C-seq sites (**F**) is significantly higher than the random expectation (mean values are indicated).

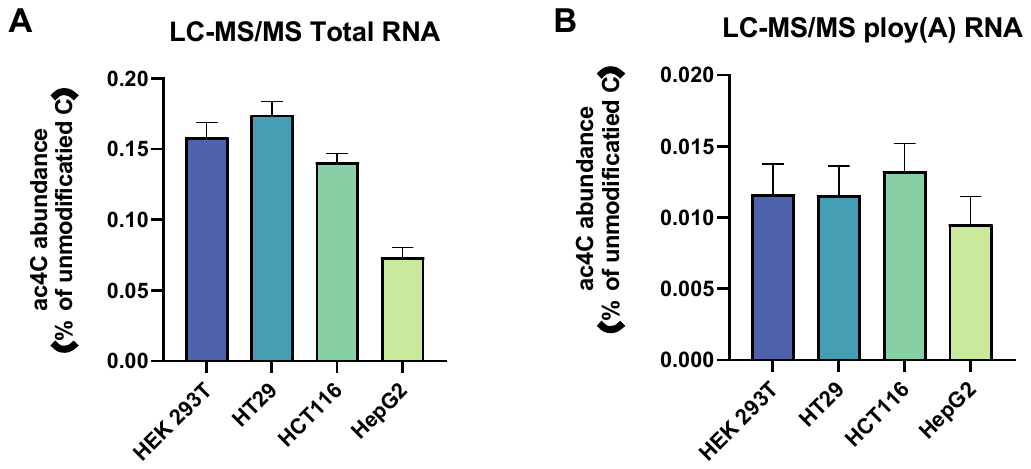

**Figure S4**. **LC-MS/MS analysis of four cell lines.** LC-MS/MS of **(A)** total RNA and **(B)** poly(A) RNA on HEK293T, HT29, HCT116, and HepG2, respectively. Mean ±SD, n=2 biological replicates x 2 technical replicates.
