## Supplementary Methods for "OpenAc4C: A gateway to decode the landscape, regulation and pathogenesis of N4-acetylcytidine (ac^4^C) epitranscriptome"

**Human cell culture and isolation of poly(A)^+^ RNA**

HEK293T/HT29/HCT116/HepG2 cells) were cultured in DMEM medium (Corning) containing 10% FBS (Cellmax) and 1% penicillin-streptomycin (Corning) under conditions of 37°C and 5% CO_2_. Cells were washed once by PBS and collected for the isolation of total RNA using TRIzol reagent (Magen). Poly(A)^+^ RNA was isolated from total RNA using oligo(dT)_25_ Dynabeads (Thermo Fisher Scientific). Total RNA and Poly(A)^+^ RNA concentration was measured by a Nanodrop ultraviolet-visible light spectrophotometer (Thermo) and Qubit 4 (Thermo).

**Determination of ac^4^C in Total RNA and Poly(A)^+^ RNA using LC-MS/MS analysis**

200 ng Total RNA or Poly(A)^+^ RNA was digested by the combination of 1 U of nuclease P1 (Wako) in 100 mM ammonium acetate for 4 hours at 42°C in 20 μl water solution, and later add 1 U of shrimp alkaline phosphatase (rSAP, NEB) and 2.5μl 10×Cutsmart buffer at 37°C for 4 h. Digested samples were filtered through 0.22-mm syringe filters before ultrahigh-performance LC–MS/MS analysis. The nucleosides were separated by a ultrahigh-performance liquid chromatographer (Shimadzu) equipped with a ZORBAX SB-Aq column (Agilent) and detected by MS/MS using a Triple Quad 5500 (AB SCIEX) mass spectrometer in positive ion mode by multiple reaction monitoring. Nucleosides were quantified using the nucleoside-to-base ion mass transitions of m/z 268.0 to 136.0 (A), m/z 245.0 to 113.1 (U), m/z 244.0 to 112.1 (C), m/z 284.0 to 152.0 (G), m/z 286.1 to 154.1 (ac^4^C). Standard curves were generated by running a concentration series of pure commercial nucleosides. Concentrations of nucleosides in samples were calculated by fitting the signal intensities to the standard curves. Following the established convention in the epitranscriptomics field [1, 2], the relative abundance of ac^4^C was expressed as the molar ratio of the modified nucleoside to its unmodified counterpart (ac^4^C/C). This ratio is calculated using the formula: % of C = (molar amount of ac^4^C / molar amount of C) × 100%.

**NaBH**_4_ **treatment and RedaC:T-seq library preparation**

SK-HEP-1 cell lines were obtained from Beijing Biocytogen and Saiweier Biotechnology, all of which underwent STR authentication to ensure correct cell origin and PCR testing to confirm the absence of mycoplasma contamination. The treatment group of NAT10-knockdown was performed by using NAT10-siRNA. 10 μg of total RNA was extracted to remove rRNA and subsequently subjected to 2 mM sodium borohydride treatment (at 55°C for 1 hour) before constructing cDNA libraries and sequencing. Other detailed process of both sample preparation of control group and treatment group was conducted by following previously reported methods for RedaC:T-seq [3].

**Quantification and statistical analysis**

The raw reads files were obtained from sequencing facility. Adapter sequences of fastq files were trimmed by using TrimGalore (v1.15) (<https://github.com/FelixKrueger/TrimGalore>) with default parameters. Then the fastq files were aligned to Human reference genome (hg38) by using STAR (v2.7.6a) [4]. And then merge and index the alignments as desired by using samtools (v1.7) [5]. For the variant analysis, run the mpileup command and pipe to the mpileup2readcounts script. Parsing the output file for tidier results for comparing mismatch rates. The parsing script was obtained from Github: (<https://github.com/dsturg/RedaCT-Seq>). Downstream analysis was conducted in R. The detailed analysis steps are presented in <https://note.youdao.com/s/7n4hTT18>.

**Identifying ONT-based ac^4^C sites using DirectRM pipeline**

To identify and quantify RNA modifications from native electronic signals, DirectRM analytical pipeline was utilized. Raw nanopore direct RNA sequencing data, acquired natively in Pod5 format for RNA004 chemistry or converted from fast5 format for RNA002 datasets, were first subjected to base-calling and reference mapping using Dorado (v0.6.2, <https://github.com/nanoporetech/dorado>). The resulting sequence alignments were subsequently compressed, sorted, and indexed utilizing Samtools (v1.7) [1] to generate standard BAM files. Next, re-squiggling was performed using the ONT Remora algorithm (v3.3, <https://github.com/nanoporetech/remora>) to precisely synchronize the raw electrical signal events with the base-called sequences. Following this data preprocessing phase, the core DirectRM framework was executed to systematically extract molecule-level features using a 9-mer sliding window. The pipeline initially deployed its *de novo* detection module to screen for candidate modified kmers based on observed signal deviations and base-calling error profiles. These candidate regions were then automatically processed through the integrated multi-label neural network to determine the specific modification type for ac^4^C, alongside its exact transcriptomic coordinate. Modification events were positively called when the predictive probability exceeded a 0.5 threshold, and any conflicting positional estimations from overlapping windows were resolved via weighted averaging. Finally, the pipeline collated these single-molecule predictions to estimate aggregated site-level or transcript-level modification rates. Putative ac^4^C sites files from multiple samples were imported into R for data filtering and integration.

**Web-based analysis tools**

The ac4CFinder and ac4CSNPer were built upon our previously developed deep-learning model using a weakly supervised learning approach (Huang et al, *Bioinformatics*, 2021) [6], which take labels at the sequence level (rather than a nucleotide level) as input and predicts the sub-regions that are most likely to contain the ac^4^C RNA modification. The pseudo ac^4^C sites (negative dataset) in our analysis were selected randomly from all available cytosines located within the same transcripts as the experimentally identified base-resolution ac^4^C sites. Briefly, the prediction process is divided into several sub-sections. Firstly, using a multi-instance learning approach, the entire RNA sequence is treated as a ‘bag’, which is split into multiple RNA instances using a 40-nt fixed-length sliding window. One-hot encoding is applied to extract sequence features from these RNA instances. The RNA instances are then fed into the model, where network weights are shared, to output instance-level features. Gated attention is used as the scoring function to obtain bag-level probabilities from these instance-level features. The bag-level feature of the entire RNA sequence (the final probability score) is determined by the weighted summation of the multiple instance features. In application, the prediction model accepts multiple standard FASTA files as input and treats each entire input sequence as a ‘bag’, reporting its bag-level label as the ac^4^C probability. The recommended sequence length for accurate prediction is around 150 nt, with a minimum of 41 nt.
